# Maternal vitamin D deficiency increases the risk of obesity in male mice offspring by affecting the immune response

**DOI:** 10.1101/2020.03.23.004721

**Authors:** Pei Li, Ping Li, Yuanlin Liu, Weijiang Liu, Lanlan Zha, Xiaoyu Chen, Rongxiu Zheng, Kemin Qi, Yi zhang

## Abstract

Recently, many epidemiological and animal studies have indicated that obesity have their origin in the early stages of life including the inappropriate balance of some nutrients, the objective of this study is to determine the risk of obesity in male mice offspring as a consequence of maternal VD deficiency-mediated disordering of the immune response. Four-week-old C57BL/6J female mice were fed VD-deficient or normal reproductive diets during pregnancy and lactation. Their male offspring were weighted and euthanized after being fed control and high-fat diets (HFD) for 16 weeks starting at the weaning. The serum was collected for biochemical analyses. Epididymal (eWAT) and inguinal (iWAT) white adipose tissues were excised for histological examination, immunohistochemistry, gene expressions of inflammatory factors, and for determining the proportions of immune cells by flow cytometry. Insufficient maternal VD intake exacerbated the development of obesity both in non-obese and obese male offspring as evidenced by larger adipose cells and abnormal glucose and lipid metabolisms. Also, the expression of proinflammatory cytokine genes was increased and that of anti-inflammatory cytokines was decreased in maternal VD-deficient groups in the eWAT and/or iWAT. This was accompanied by higher levels of TNF-α or/and INF-β, and lower levels of IL-4 and IL-10. Insufficient maternal VD intake was also observed to induce a shift in the profiles of immune cells in the eWAT and/or iWAT, resulting in increased percentages of M1 macrophage, ATDCs, and CD4^+^ and CD8^+^ T cells, but caused a significant decrease in the percentage of M2 macrophages, both in non-obese and obese male offspring. All these changes in the immune cell profile were more obvious in the eWAT than in the iWAT. These results indicated that insufficient maternal VD intake promoted the development of obesity in male offspring by modulating the immune cell populations and causing a polarization in the adipose depots.

**Importance:** Evidence in this study has indicated that insufficient maternal VD intake promotes the development of obesity in the male offspring by modulating the recruitment of immune cell populations and their polarization as well as the expression and secretion of proinflammatory adipokines in the adipose depots in a weight-independent manner, which is more obvious in eWAT than that in the iWAT.

## Introduction

The prevalence of overweight and obesity has reached epidemic proportions throughout the world, and is associated with increased morbidity and mortality, contributing to the burden of chronic diseases (1–3). However, the mechanisms underlying obesity remain unclear. There is, therefore, an urgent need for effective intervention strategies. During the past decade, the state of chronic low-grade systemic inflammation caused by inflation of adipose tissue macrophages (ATMs), followed by increased production of pro-inflammatory cytokines, such as TNF-α, IL-6, and decreased contents of anti-inflammatory factors, including IL-10, has been recognized as one of the factors in the pathogenesis of obesity (4–6). It is well established that the crosstalk between adipocyte hypertrophy and inflammation can exacerbate chronic inflammation (7,8). Macrophages (M) make up the majority of leukocytes infiltrating the adipose tissue (9). The infiltration of M1, the pro-inflammatory ATMs, represents a key event influencing the adipose tissue dysfunction. With increasing adiposity, the polarization status of anti-inflammatory M2 macrophages switches to a more pro-inflammatory M1 state so that these macrophages jointly promote the development of obesity^10^. Obesity is associated with adipocyte hypertrophy, which can cause rupturing of adipocytes and result in increased local accumulation of inflammatory cells, including M1 and M2 macrophages and Th1/Th2 cells, as well as in the altered production of adipokines (11). The increase in the numbers of M1 can elicit effector functions through a surplus production of pro-inflammatory cytokines and promotes insulin resistance in adipocytes (12). The decreased secretion of inflammatory adipokines from obese adipose tissue can decrease the ongoing recruitment and polarization of macrophages to exacerbate chronic inflammation in obesity (13,14).

Vitamin D (VD) comprises an important group of fat-soluble seco-sterols required for bone growth and calcium homeostasis, as well as for its role as an enzyme agonist. There is a strong association between VD status and obesity, with low VD levels being highly prevalent in obese people (15,16). Achieving dietary reference intake through VD supplements could reduce body weight by influencing calcium absorption (17), parathyroid hormone expression (18), phosphate metabolism (19), growth-plate function (20), and regulation of inflammatory reactions (21). *In vitro* experiments have confirmed that VD has immunoregulatory effects and can reduce adipocyte inflammation, which may participate in the reduction of adipose tissue macrophage infiltration in the context of obesity-associated low-grade inflammation (22). Besides, animal experiments have also shown that dietary insufficiency of VD exacerbates high-fat diet-induced gain in body weight, adipose tissue expansion and macrophage infiltration, and inflammation (23). However, the specific mechanisms for such effects remain unclear.

Recently, many epidemiological and animal studies have indicated that overweight and obesity have their origin in the early stages of life; for example, an inappropriate balance of some nutrients during the early infancy or fetal life, can permanently alter the properties of fatness (24). Under these conditions, timely and accurate nutritional intervention during the perinatal period is likely to be more effective than the interventions for improvement of metabolic health in the later part of life (25). Vitamin D deficiency is prevalent among all age groups, and especially in pregnant women who require higher amounts of VD for their own metabolism as well as for their baby (26). In an epidemiological research, low VD status was found in about 69% of the pregnant women in China (27). Despite the known effects of maternal VD deficiency on the obesity of the offspring, clinical and animal studies designed to evaluate the effect on body fat mass in the offspring have produced mixed results owing to several methodological limitations (28–31). Moreover, only a few studies have reported on the possible mechanisms underlying the effects of maternal VD deficiency on the body weight of the offspring expect for its effects on adipocyte proliferation (30). Therefore, in the present study, we investigated the effects of insufficient VD intake through diet during maternal pregnancy and lactation on the body weight gain in the male offspring and on the immune response using a high-fat diet-induced obese mouse model.

## Results

### Maternal and offspring VD status

The concentrations of 25(OH)D3 were assessed in five mice from each group of mother and 21-day-old male offspring (Supplement Figure 2) to confirm that the contents of vitamin D *in vivo* were consistent with the dietary regimen [Mother: 134.14 μg/L (VD-C), 43.87 μg/L (VD-D); 21-day-old male offspring: 251.66 μg/L (VD-C), 167.78 μg/L (VD-D)]. As shown in Table 1, no differences existed in the levels of 25(OH)D3 in either non-obese (VD-C and VD-D) or obese (VD-C-HFD and VD-D-HFD) male offspring born to females in the maternal VD-sufficient or-deficient groups. However, the levels were lower in all the obese groups than that in the non-obese groups (*P*<0.05).

**Table 1.**
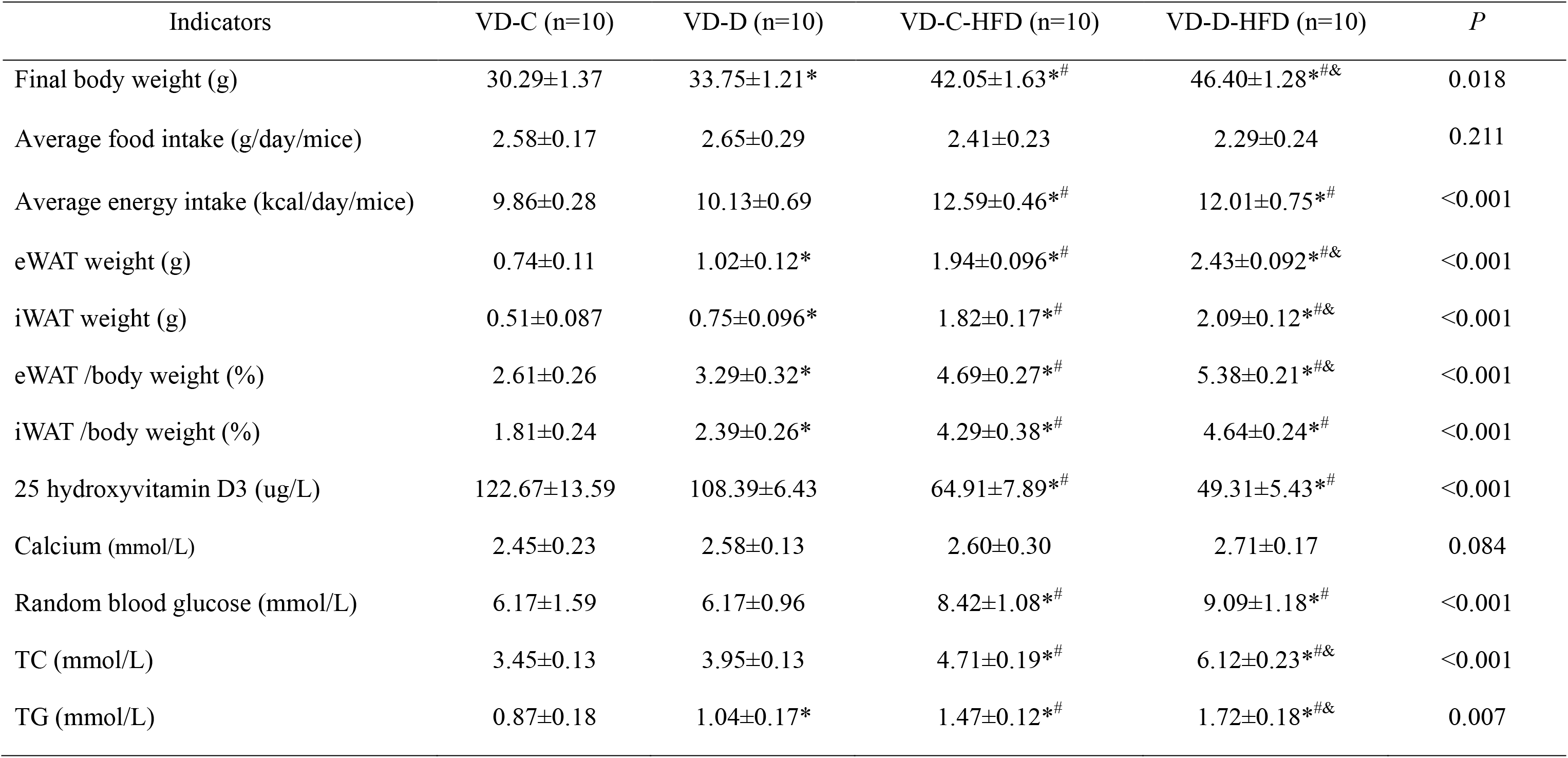
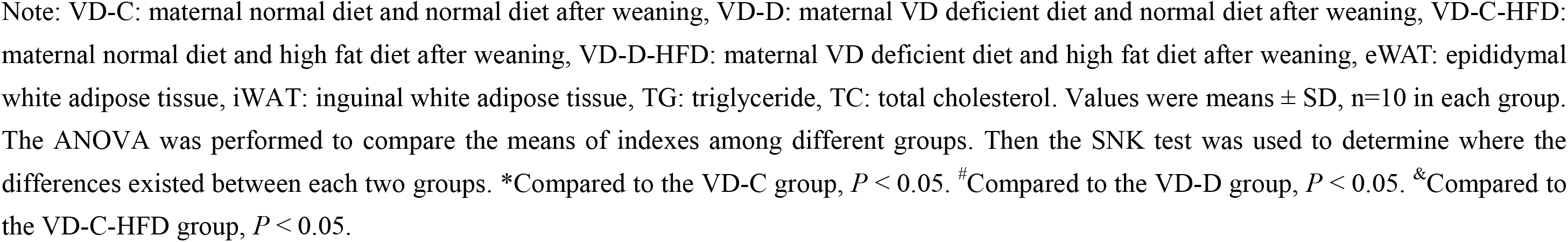
Effects of maternal vitamin D status on the metabolic characteristics among the male offspring

| Indicators | VD-C (n=10) | VD-D (n=10) | VD-C-HFD (n=10) | VD-D-HFD (n=10) | *P* |
| --- | --- | --- | --- | --- | --- |
| Final body weight (g) | 30.29±1.37 | 33.75±1.21* | 42.05±1.63*<sup>#</sup> | 46.40±1.28*<sup>#&</sup> | 0.018 |
| Average food intake (g/day/mice) | 2.58±0.17 | 2.65±0.29 | 2.41±0.23 | 2.29±0.24 | 0.211 |
| Average energy intake (kcal/day/mice) | 9.86±0.28 | 10.13±0.69 | 12.59±0.46*<sup>#</sup> | 12.01±0.75*<sup>#</sup> | <0.001 |
| eWAT weight (g) | 0.74±0.11 | 1.02±0.12* | 1.94±0.096*<sup>#</sup> | 2.43±0.092*<sup>#&</sup> | <0.001 |
| iWAT weight (g) | 0.51±0.087 | 0.75±0.096* | 1.82±0.17*<sup>#</sup> | 2.09±0.12*<sup>#&</sup> | <0.001 |
| eWAT /body weight (%) | 2.61±0.26 | 3.29±0.32* | 4.69±0.27*<sup>#</sup> | 5.38±0.21*<sup>#&</sup> | <0.001 |
| iWAT /body weight (%) | 1.81±0.24 | 2.39±0.26* | 4.29±0.38*<sup>#</sup> | 4.64±0.24*<sup>#</sup> | <0.001 |
| 25 hydroxyvitamin D3 (ug/L) | 122.67±13.59 | 108.39±6.43 | 64.91±7.89*<sup>#</sup> | 49.31±5.43*<sup>#</sup> | <0.001 |
| Calcium (mmol/L) | 2.45±0.23 | 2.58±0.13 | 2.60±0.30 | 2.71±0.17 | 0.084 |
| Random blood glucose (mmol/L) | 6.17±1.59 | 6.17±0.96 | 8.42±1.08*<sup>#</sup> | 9.09±1.18*<sup>#</sup> | <0.001 |
| TC (mmol/L) | 3.45±0.13 | 3.95±0.13 | 4.71±0.19*<sup>#</sup> | 6.12±0.23*<sup>#&</sup> | <0.001 |
| TG (mmol/L) | 0.87±0.18 | 1.04±0.17* | 1.47±0.12*<sup>#</sup> | 1.72±0.18*<sup>#&</sup> | 0.007 |

Note: VD-C: maternal normal diet and normal diet after weaning, VD-D: maternal VD deficient diet and normal diet after weaning, VD-C-HFD: maternal normal diet and high fat diet after weaning, VD-D-HFD: maternal VD deficient diet and high fat diet after weaning, eWAT: epididymal white adipose tissue, iWAT: inguinal white adipose tissue, TG: triglyceride, TC: total cholesterol. Values were means ± SD, n=10 in each group. The ANOVA was performed to compare the means of indexes among different groups. Then the SNK test was used to determine where the differences existed between each two groups. \*Compared to the VD-C group, $P < 0.05$ . #Compared to the VD-D group, $P < 0.05$ . &Compared to the VD-C-HFD group, $P < 0.05$ .

### Body and adipose tissue weight

As shown in Table 1 and Figure 1, there were maternal effects of VD intake on the body and adipose tissue weight, both in the non-obese and obese male offspring. The body, eWAT, and iWAT weights and the percentages of eWAT and iWAT were prominently higher in mice from dams of the VD-D group, than in those from the VD-C group; the former also had larger adipose cells in the eWAT (Figure 1C and 1D) and iWAT (Figure 1E and 1F) (*P*<0.05). In contrast, among the obese offspring, the body, eWAT, and iWAT weights, and the percentages of eWAT were higher and the volume of fat cells in eWAT was larger in the VD-D-HFD group than in the VD-C-HFD group (*P*<0.05).

**Figure 1.**
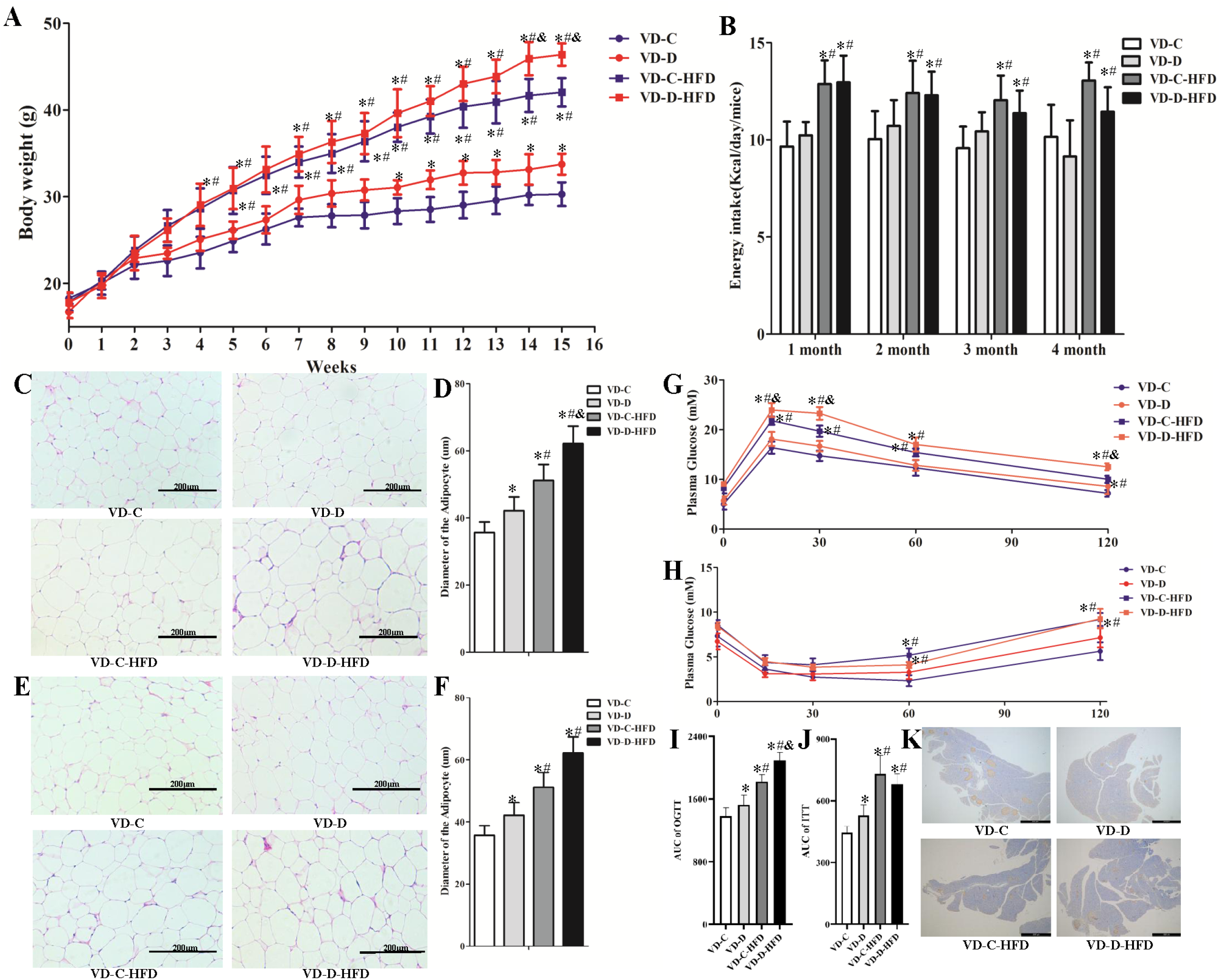
Body weight, glucose, and lipid metabolism-related indicators in response to different maternal vitamin D intake in the non-obese and obese male offspring. The body weights in the maternal vitamin D-deficient groups were found to be significantly increased compared to that in the control group both in the non-obese (VD-D vs. VD-D) and obese male offspring (VD-D-HFD vs. VD-C-HFD) (A), and were accompanied with larger adipose cells in the eWAT (C and D) and iWAT (E and F), decreased insulin secretion (H, J, and K), and higher glucose levels in the serum (G and I), with no differences in the energy intake (B). n = 10 mice/group. VD-C: maternal normal diet and normal diet after weaning, VD-D: maternal VD deficient diet and normal diet after weaning, VD-C-HFD: maternal normal diet and high-fat diet after weaning, VD-D-HFD: maternal VD-deficient diet and high-fat diet after weaning, eWAT: epididymal white adipose tissue, iWAT: inguinal white adipose tissue. n = 10 in each group. * *P*<0.05, compared to the VD-C. ^#^ *P*<0.05, compared to the VD-D. ^&^ *P*<0.05, compared to the VD-C-HFD.

### Glucose and lipid metabolism-related indicators in the serum

The baseline fasting glucose concentrations were not affected by the different maternal VD intake during pregnancy and lactation both in non-obese and obese male offspring, whereas higher glucose levels were observed in the serum in the obese groups (VD-C-HFD and VD-D-HFD) than in the non-obese groups (VD-C and VD-D) (*P*<0.05) (Table 1). Likewise, the circulating glucose response to the glucose load, as indicated by OGTT, ITT, and related by the area under the curve (AUC) (Figure 1G 1H, 1I and 1J), showed that the glucose levels in the maternal VD-deficient group increased both in the non-obese (VD-D vs. VD-C) and obese (VD-D-HFD vs. VD-C-HFD) male offspring after intraperitoneal glucose administration (*P*<0.05). However, the results of ITT showed that the offspring from maternal VD-sufficient mice did not show improved insulin tolerance. Besides, the results of immunohistochemistry showed that the number of insulin-positive cells and insulin secretion decreased significantly in the maternal VD-deficient group both in the non-obese (VD-D vs. VD-C) and obese (VD-D-HFD vs. VD-C-HFD) male offspring (*P*<0.05).

The concentrations of lipid metabolism-related indicators, namely TG and TC, are presented in Table 1. The concentrations of TG and TC in the obese (VD-D-HFD and VD-C-HFD) groups were higher than those in the non-obese (VD-D and VD-C) groups. Furthermore, comparison between the two similar groups showed that the contents of TG increased in the VD-D group compared to that in the VD-C group, whereas among the obese groups, the concentrations of TG and TC were higher in the VD-D-HFD group than in the VD-C-HFD group (*P*<0.05).

### Infiltration and percentage of immune cells in the eWAT and iWAT

The infiltration of inflammatory factors in eWAT and iWAT, shown by F4/80 in Figure 2C and D, was more prominent in the obese (VD-D-HFD vs. VD-D and VD-C-HFD vs. VD-C) groups, whereas more inflammatory factors were present in the eWAT in maternal VD-deficient group both in the non-obese (VD-D vs. VD-C) and obese (VD-D-HFD vs. VD-C-HFD) male offspring. However, in the iWAT, these factors were consistently found in the non-obese male offspring (VD-D vs. VD-C).

**Figure 2.**
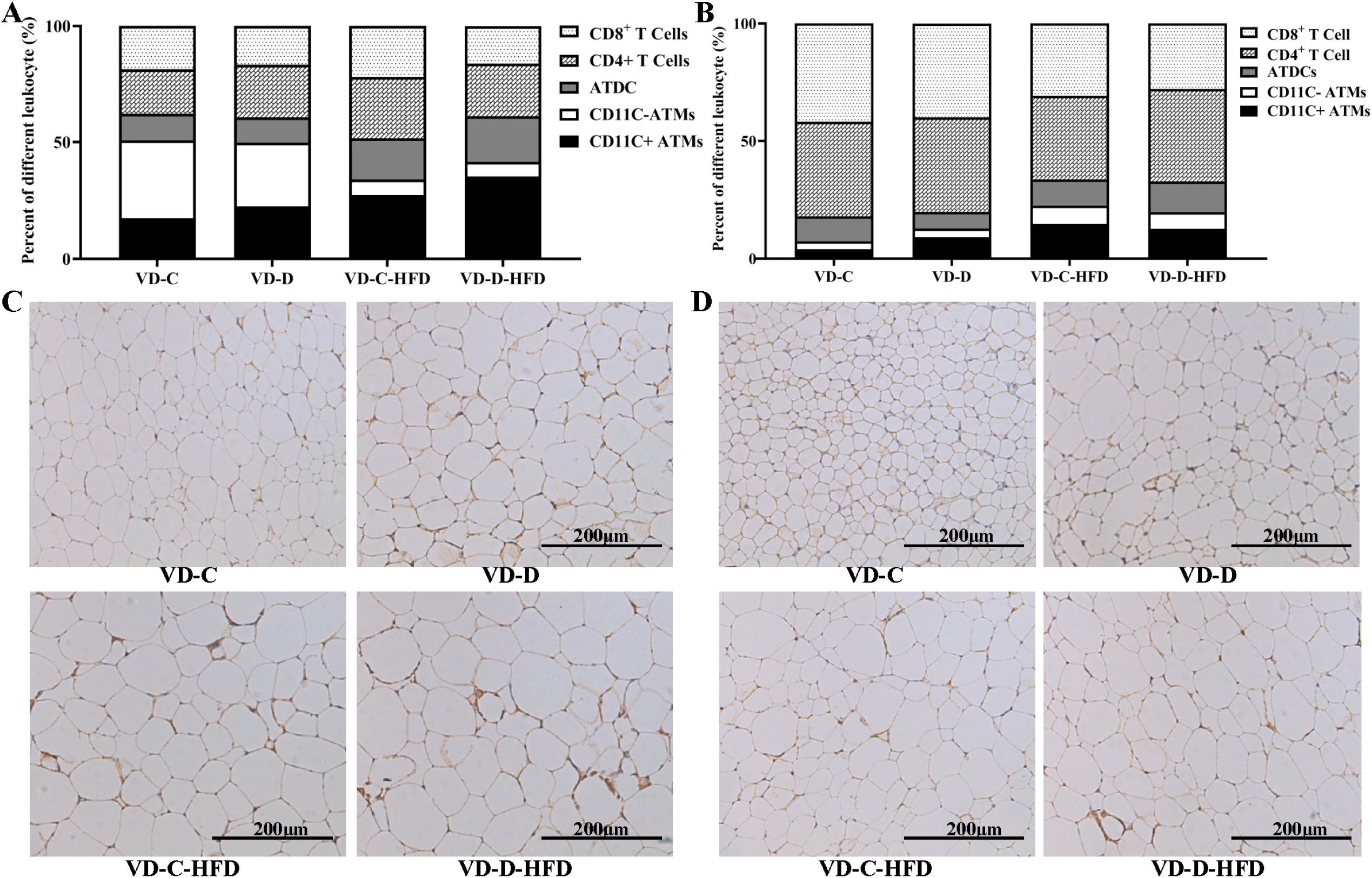
Infiltration and percentage of immune cells in the eWAT and iWAT in response to different maternal vitamin D intake in the non-obese and obese male offspring. The infiltration and percentage of immune cells was affected by the different maternal vitamin D intake both in the non-obese and obese male offspring in the eWAT (A and C) and iWAT (B and D). VD-C: maternal normal diet and normal diet after weaning, VD-D: maternal VD-deficient diet and normal diet after weaning, VD-C-HFD: maternal normal diet and high-fat diet after weaning, VD-D-HFD: maternal VD-deficient diet and high-fat diet after weaning.

The percentages of immune cells, including CD11C^+^ ATMs, CD11C^−^ ATMs, ATDCs, CD4^+^ T cells, and CD8^+^ T cells, in the eWAT and iWAT are presented in Figure 2A and 2B. In the eWAT, the percentages of CD11C^+^ ATMs, ATDCs, and CD4^+^T cells were higher, and that of CD11C^−^ ATMs were lower in the obese (VD-D-HFD and VD-C-HFD) groups than in the non-obese groups (VD-D and VD-C) (*P*<0.05). Further comparison between the two similar groups showed that maternal VD deficiency during pregnancy and lactation could increase the percentages of CD11C^+^ ATMs, ATDCs, and CD4^+^ T cells and decrease the percentages of CD11C^−^ ATMs and CD8^+^ T cells both in the non-obese (VD-D vs. VD-C) and obese (VD-D-HFD vs. VD-C-HFD) male offspring (*P*<0.05). However, in the iWAT, the percentages of CD11C^+^ ATMs, CD11C^−^ ATMs, and ATDCs were higher, and that of CD8^+^ T cells was lower in the obese (VD-D-HFD and VD-C-HFD) groups than in the non-obese (VD-D and VD-C) groups (*P*<0.05). Further comparison among the two similar groups showed that maternal VD deficiency during pregnancy and lactation could increase the percentages of CD11C^+^ ATMs and decrease the contents of CD8^+^ T cells both in the non-obese (VD-D vs. VD-C) and obese (VD-D-HFD vs. VD-C-HFD) male offspring (*P*<0.05).

### Population of ATM, ADTC, and T cells in the eWAT and iWAT

Within the adipose tissues, CD45^+^CD64^+^ cells in the SVFs were identified as the ATMs, which were then further characterized as CD11C^+^ ATMs (M1 ATMs, M1 macrophages) and CD11C^−^ ATMs (M2 ATMs, M2 macrophages). The adipose tissue dendritic cells 187 (ATDCs) were identified as CD45^+^CD64^−^CD11C^+^. The T cells were identified as CD45^+^CD3^+^, and were further categorized as CD4^+^ and CD8^+^ T cells.

In the eWAT (see Figure 3), M1 ATMs, M1 ratio (M1/M2), ATDCs, CD4^+^, and CD8^+^ T cells increased, and M2 ATMs and M2 ratio (M2/M1) decreased with the feeding of HFD (VD-C-HFD vs. VD-C and VD-D-HFD vs. VD-D) (*P*<0.05), with no changes in the ratio of CD4^+^/CD8^+^ T cells. Maternal VD deficiency during pregnancy and lactation led to significant increases in M1 ATMs, M1 ratio (M1/M2), and ATDCs, and reduced the M2/M1 ratio both in the non-obese (VD-D vs. VD-C) and obese (VD-D-HFD vs. VD-C-HFD) groups (*P*<0.05). Besides, maternal VD deficiency caused a weight-independent increase in CD4^+^ T cells in the non-obese (VD-D vs. VD-C) groups (*P*<0.05). In the iWAT, the M1 ATMs, M2 ATMs, ATDCs, CD4^+^ T cells, M1/M2 ratio, and CD4^+^/CD8^+^ T cells increased, and CD8^+^/CD4^+^ T cells decreased with HFD feeding (*P*<0.05), but there was no effect on the proportion of CD8^+^/CD4^+^ T cells (Figure 4). In contrast, there was a significant weight-independent upregulation in the proportion of ATDCs both in the non-obese (VD-D vs. VD-C) and obese (VD-D-HFD vs. VD-C-HFD) groups (*P*<0.05). Within the control groups, it was observed that M1 ATMs, M2 ATMs, CD4^+^ T cells, CD8^+^ T cells, and M1/M2 ratio were higher in the VD-D group than in the VD-C group (*P*<0.05), which were not matched by all types of immune cells, only leaving higher percentage of CD4^+^ T cells (CD4^+^/CD8^+^) and lower percentage of CD8^+^ T cells (CD8^+^/CD4^+^) in the VD-D-HFD group than those in the VD-C-HFD group (*P*<0.05).

**Figure 3.**
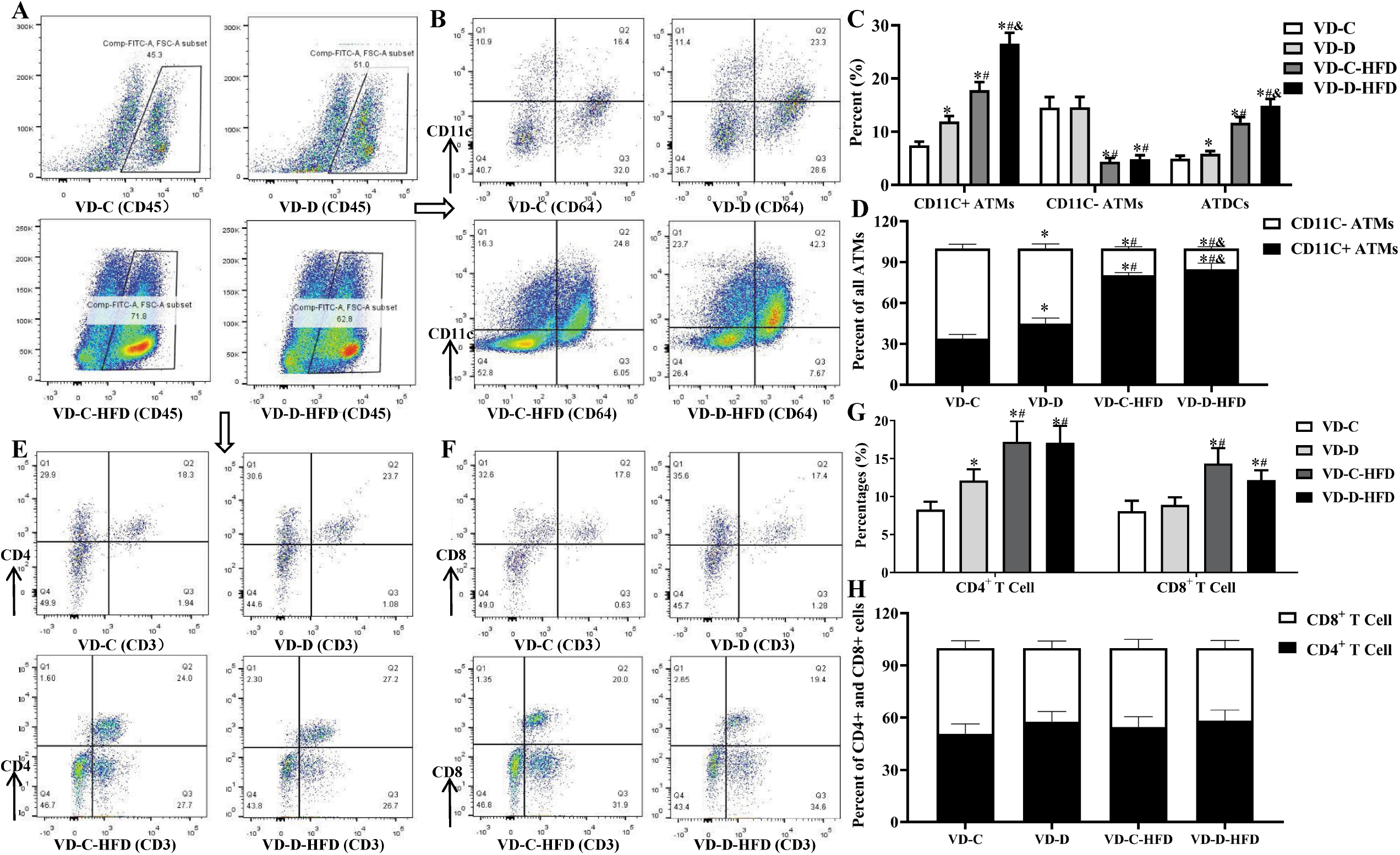
Population of ATM, ADTC, and T cells in the eWAT in response to different maternal vitamin D intake in the non-obese and obese male offspring. Maternal VD-deficient intake increased the population of CD11C^+^ATMs (A, B, and C), ATDCs (A, B, and C), CD4^+^T cells (E, F, and G), CD8^+^ T cells (E, F, and G), the percentage of CD11C^+^ ATMs (D), and decreased the proportion of CD11C^−^ ATMs (A, B, and C) and the percentage of CD11C^−^ ATMs (D), with no differences in the ratio of CD4^+^/CD8^+^ T cells (H) both in the non-obese (VD-D vs. VD-D) and/or obese (VD-D-HFD vs. VD-C-HFD) male offspring. VD-C: maternal normal diet and normal diet after weaning, VD-D: maternal VD deficient diet and normal diet after weaning, VD-C-HFD: maternal normal diet and high-fat diet after weaning, VD-D-HFD: maternal VD-deficient diet and high-fat diet after weaning, eWAT: epididymal white adipose tissue. * *P*<0.05, compared to the VD-C. ^#^ *P*<0.05, compared to the VD-D. ^&^ *P*<0.05, compared to the VD-C-HFD.

**Figure 4.**
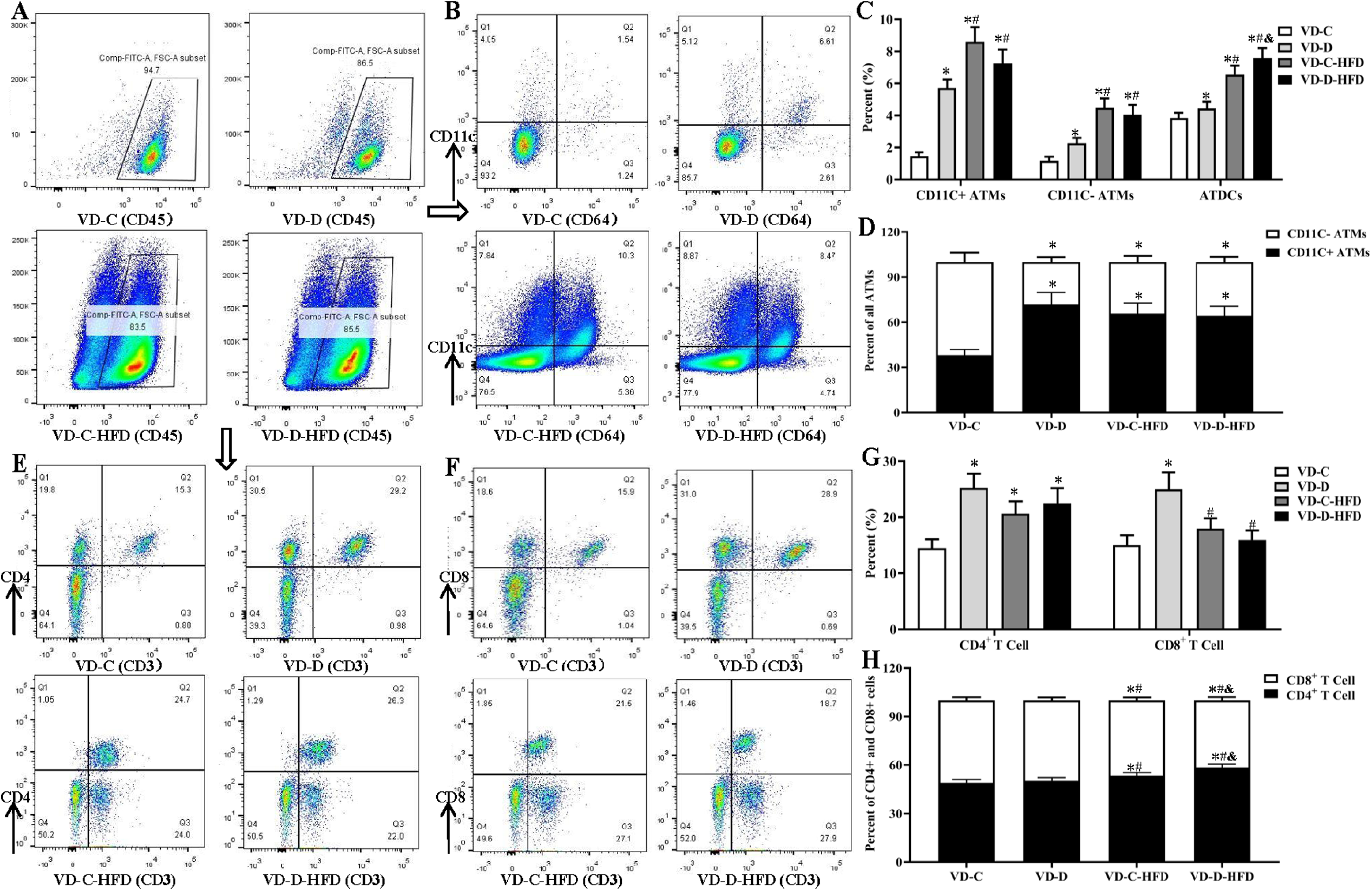
Population of ATM, ADTC, and T cells in the iWAT in response to different maternal vitamin D intake in the non-obese and obese male offspring. Maternal VD-deficient intake increased the population of CD11C^+^ ATMs (A, B, and C), CD11C^−^ ATMs (A, B, and C), ATDCs (A, B, and C), CD4^+^ T cells (E, F, and G), CD8^+^ T cells (E, F, and G), the percentage of CD11C^+^ATMs (D) and CD4^+^ T (H) in the non-obese groups (VD-D vs. VD-D). In the obese groups, the population of ATDCs (A, B, and C) and the percentage of CD4^+^ T cells (H) were higher in VD-D-HFD than in the VD-C-HFD group. VD-C: maternal normal diet and normal diet after weaning, VD-D: maternal VD-deficient diet and normal diet after weaning, VD-C-HFD: maternal normal diet and high-fat diet after weaning, VD-D-HFD: maternal VD-deficient diet and high-fat diet after weaning, iWAT: inguinal white adipose tissue. * *P*<0.05, compared to the VD-C. ^#^ *P*<0.05, compared to the VD-D. ^&^ *P*<0.05, compared to the VD-C-HFD, *P*<0.05.

### Gene expression profiles of proinflammatory and inflammasome cytokines in the eWAT and iWAT

To examine the effects of different maternal vitamin D intake during pregnancy and lactation on the expression of proinflammatory (INOS, IL-1β, TNF-α, INF-γ, and IL-6) and anti-inflammatory (Arg-1, IL-4, and IL-10) cytokine genes in the eWAT and iWAT, we analyzed the mRNA levels of these genes. Compared with the VD-C group, the expression levels of all the proinflammatory and anti-inflammatory cytokines were significantly increased both in the eWAT and iWAT of VD-D, VD-C-HFD, and VD-D-HFD mice, whereas the expression of Arg-1 was apparently reduced (Figure 5, *P*<0.05). Moreover, in the obese mice, the mRNA levels of INOS, IL-1β, TNF-α, INF-γ, and IL-6 were higher and those of Arg-1, IL-4, and IL-10 were lower in the VD-D-HFD group than in the VD-C-HFD group in the eWAT (*P*<0.05). Consistently, the levels of INF-γ were significantly increased, whereas those of Arg-1 were decreased in the iWAT of VD-D-HFD mice compared to the respective levels in the VD-C-HFD mice (*P*<0.05). However, the effects on the other cytokines in the iWAT were not significantly different. Taken together, these results indicate that maternal VD status could affect the expression of some proinflammatory and inflammasome cytokine genes in the eWAT and iWAT, and that the differences were greater in the eWAT.

**Figure 5.**
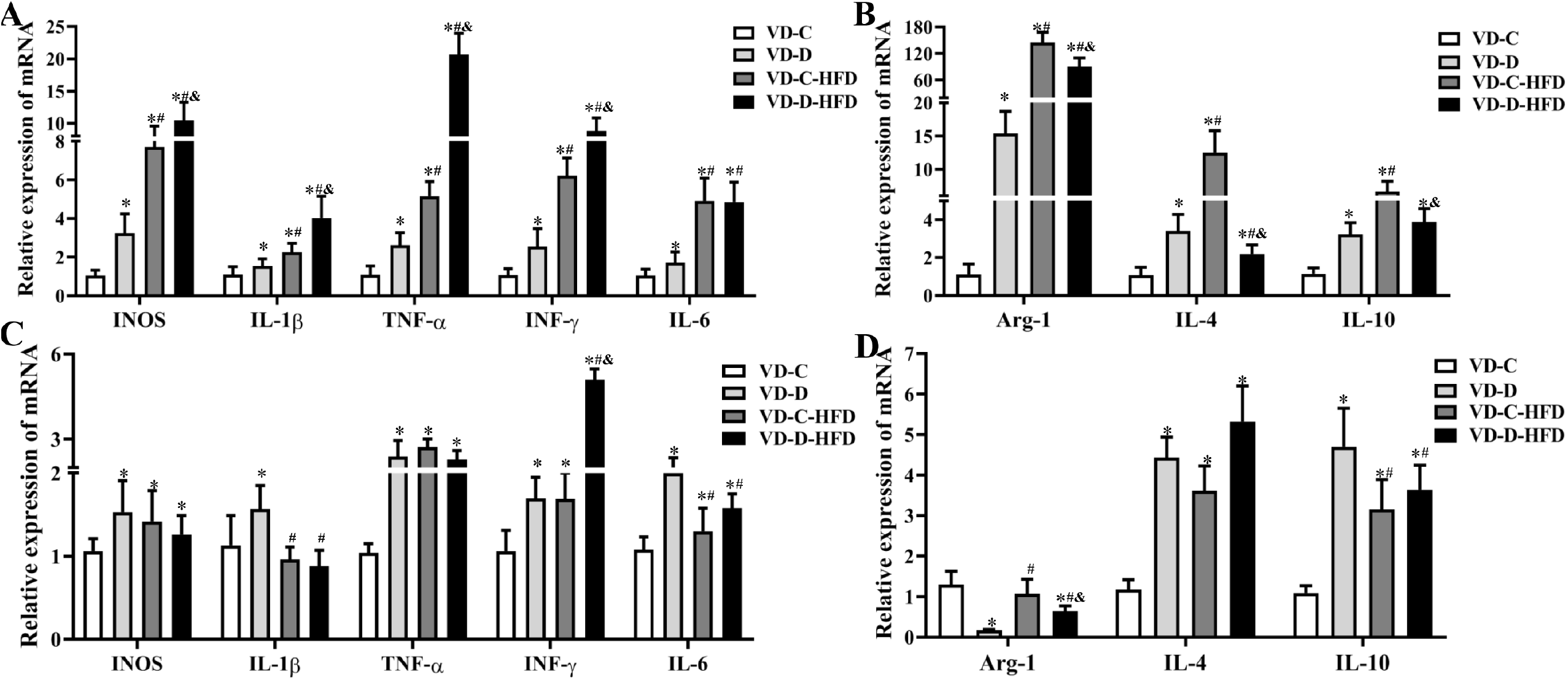
Expression levels of proinflammatory and inflammasome cytokine genes in the eWAT and iWAT. Maternal VD-deficient intake increased the expression levels of inflammasome cytokine genes, namely INOS, IL-1β, TNFα, INFβ, and IL-6 in the eWAT both in the non-obese (VD-D vs. VD-D) and obese male offspring (VD-D-HFD vs. VD-C-HFD) (A), whereas the expression levels of genes encoding the proinflammatory cytokines, including Arg-1, IL-4, and IL-10, were higher in the VD-D group than in the VD-C group, and were lower in VD-D-HFDgroup than in the VD-HFD group in the eWAT (B). In the iWAT, maternal VD-deficient intake increased the levels of genes encoding the cytokines, INOS, IL-1β, TNFα, INFβ, IL-6, IL-4, and IL-10, in the non-obese (VD-D vs. VD-C) male offspring (C, D), whereas in the obese group, the expression levels of the INFβ gene were higher and those of the Arg-1 gene were lower in the VD-D-HFD group than in the VD-C-HFD group. VD-C: maternal normal diet and normal diet after weaning, VD-D: maternal VD-deficient diet and normal diet after weaning, VD-C-HFD: maternal normal diet and high-fat diet after weaning, VD-D-HFD: maternal VD-deficient diet and high fat diet after weaning, eWAT: epididymal white adipose tissue; iWAT: inguinal white adipose tissue. \**P*<0.05, compared to the VD-C. ^#^ *P*<0.05, compared to the VD-D. ^&^ *P*<0.05, compared to the VD-C-HFD.

### Levels of cytokines in the serum

The levels of serum cytokines, TNF-α, INF-γ, IL-6, IL-4, and IL-10, were determined in all the samples (Figure 6). High-fat diets resulted in a significant increase in the levels of TNF-α and INF-γ and a significant decrease in the levels of IL-4 and IL-10. Furthermore, the levels of TNF-α or/and INF-γ were higher, whereas those of IL-4 and IL-10 were lower in the maternal VD-deficient group both in the non-obese (VD-D vs. VD-C) and obese (VD-D-HFD vs. VD-C-HFD) male offspring.

**Figure 6.**
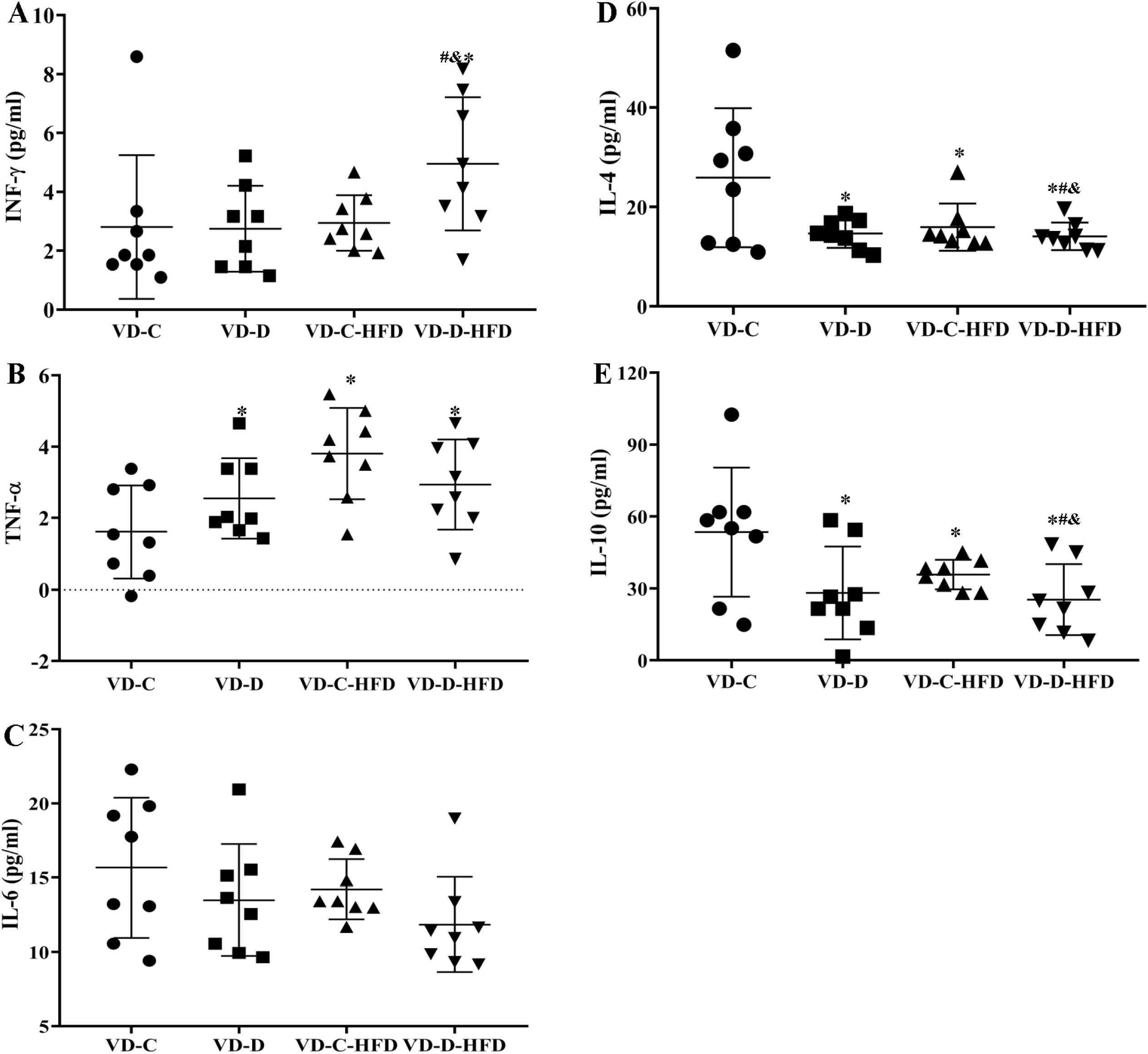
Levels of cytokines in the serum in response to different maternal vitamin D intake in the non-obese and obese male offspring. The levels of inflammasome cytokines, namely INFβ (A) and TNFα (B) in the maternal vitamin D-deficient groups were found to be significantly increased, whereas the contents of proinflammatory cytokines, IL-4 (D) and IL-10 (E), were decreased compared to that in the control group both in the non-obese (VD-D vs. VD-D) and/or obese male offspring (VD-D-HFD vs. VD-C-HFD). VD-C: maternal normal diet and normal diet after weaning, VD-D: maternal VD-deficient diet and normal diet after weaning, VD-C-HFD: maternal normal diet and high-fat diet after weaning, VD-D-HFD: maternal VD-deficient diet and high-fat diet after weaning, eWAT: epididymal white adipose tissue; iWAT: inguinal white adipose tissue. * *P*<0.05, compared to the VD-C.^#^ *P*<0.05, compared to the VD-D. ^&^ *P*<0.05, compared to the VD-C-HFD.

## Discussion

Recently, several epidemiological and experimental studies have indicated that VD supplementation is effective in reducing body weight, body-mass index, and fat mass and is associated with decreased levels of low-density lipoprotein cholesterol (32,33). In this study, we investigated whether the effects of maternal VD status during pregnancy and lactation had a long lasting adverse effect on the progress of obesity in the male offspring. We found that insufficient maternal VD intake could worsen the development of obesity both in the non-obese (VD-D vs. VD-C) and obese (VD-D-HFD vs. VD-C-HFD) male offspring, as evidenced by larger adipose cells, and abnormal glucose and lipid metabolisms in the eWAT and/or iWAT. All these results were consistent with those reported by Belenchia et al (34). All these results suggest that gestation of male murine offspring in an environment of maternal VD deficiency can lead to fetal growth restrictions, accelerated growth in early life, larger visceral body fat pads, and greater susceptibility to HFD-induced adipocyte hypertrophy. Besides, all these changes were more obvious in the eWAT than in the iWAT because obesity is mainly caused by the increase in white fat and inflammatory infiltration, and it is more complex of the cells types in eWAT to weaken the difference above.

The reasons for the observed harmful effects of insufficient maternal VD intake on the weight gain include an unhealthy trabecular bone structure (31), unreasonable colonization of intestinal flora (30), greater susceptibility to HFD-induced adipocyte hypertrophy among others (35). Adipocyte hypertrophy could lead to the alterations in immune cell populations after the onset of obesity. It is well established that the development of obesity is characterized by immune cell infiltration and low-grade inflammation in the obese adipose tissue. Macrophages make up the majority of the leukocytes infiltrating the adipose tissue, and two primary types of macrophages (ATMs) have been identified based on the expression of CD11C (36). Lean mice and humans have a preponderance of the resident population of CD11C^−^ ATMs (M2) whereas obese individuals accumulate CD11C^+^ ATMs (M1) that have a lysosomal activation phenotype (37,38). In this case, cytokines that are released by inflammatory cells infiltrating the obese adipose tissue include tumor necrosis factor-alpha (TNF-α), Interferon gamma (INF-γ), interleukin 6 (IL-6), monocyte chemoattractant protein 1 (MCP-1), and IL-1. All these molecules may act on immune cells leading to local and generalized inflammation. As expected, Elimrani et al. showed that VD supplementation reduced the severity of colitis and decreased the number of inflammation-associated colorectal tumors in C57BL/6J and diabetic mice (39). Martinez-Santibanez et al also showed that HFD-induced male obesity was associated with more weight gain and increased accumulation of CD11c^+^ ATMs, which are known to be involved in adipose tissue remodeling, in the animals in the VD-deficient group than in control group animals (40). This study indicated that appropriate intake of VD might regulate the ratio between these two types of ATMs to improve their function, helping the adipose tissue to remodel, which can contribute to adipocyte hypertrophy. We also found increased expression of proinflammatory and decreased expression of anti-inflammatory cytokines in maternal VD-deficient groups (VD-D-HFD vs. VD-C-HFD and VD-D vs. VD-C) in the eWAT and/or iWAT, accompanied by higher levels of TNF-α or/and INF-γ, and lower contents of IL-4 and IL-10. We also observed that insufficient maternal VD intake induced a shift in the immune cell profiles in the eWAT and/or iWAT, which included increased percentages of M1 macrophages both in non-obese and obese male offspring. In addition, macrophage infiltration in the adipose tissue, as demonstrated by number of CLS (F4/80-positive adipocytes surrounded by macrophages), was exacerbated by inadequate VD intake. All these immune changes were more obvious in the eWAT than in the iWAT. These results indicated that VD insufficiency-increased adipose tissue inflammation might be a consequence of diet-induced adipose expansion/ obesity together with increased inflammation in the white adipose tissue.

In obesity, the accumulation of CD4^+^ T cells and CD4^+^/CD8^+^ cells precedes macrophage infiltration to promote their recruitment to the obese adipose tissue. The increased proportion of ATDCs can also contribute to altered immune function in the adipose tissue that might aggravate the metabolic function including obesity. Recent investigations have emphasized the contributions of adaptive immune cells, especially CD4^+^ and CD8^+^ T cells that play an important role in immunity by regulating the secretion of proinflammatory cytokines, such as IFN-γ and TNF-α, to affect the progression of obesity and associated disorders (41). Moreover, the accumulation of these cells induces inflammation (42). This is further characterized by an increase in IFN-γ producing CD8^+^ T cells in the lamina propria of obese versus lean subjects, suggesting a proinflammatory shift of CD8^+^ T cells in the intestine of individuals with obesity. This proinflammatory shift, with an increased number of CD4^+^ and CD8^+^ T cells was also confirmed in another study on the colon and small intestine of persons with obesity, albeit in a smaller population(43). Additionally, we found that the proportions of CD4^+^ and CD8^+^ T cells were increased in the obese male offspring, and were also higher in the eWAT and/or iWAT of maternal VD-deficient groups (VD-D vs. VD-C), accompanied by higher levels of TNF-α or/and INF-γ. Classically, it is possible that the increased ratio of ATDCs can contribute to altered immune function in the adipose tissue that may contribute to its expansion. Zlotnikov-Klionzky et al reported that deletion of perforin positive ATDCs had a high impact on metabolic phenotype as mice lacking perforin-positive ATDCs gained weight while being maintained on a control diet, were glucose intolerant and insulin resistant, and had increased levels of lipids in the blood (44). Our observations suggest that ATDCs have the capacity to control T cell fates as antigen presenting cells and maternal VD deficiency could increase the percentages of ATDCs both in non-obese (VD-D vs. VD-C) and obese (VD-D-HFD vs. VD-C-HFD) male offspring.

In the summary, we found that insufficient maternal VD intake promotes the development of obesity in the male offspring by modulating the recruitment of immune cell populations and their polarization as well as the expression and secretion of proinflammatory adipokines in the adipose depots in a weight-independent manner.

## Materials and Methods

### Animal study

Thirty-four-weeks-old C57BL/6J female mice purchased from Charies River SPF Laboratory Animal Technology Co. Ltd. (Beijing) were housed in the laboratory Animal Center of the Institute of Military Medicine, Military Academy of Military Sciences of China under standard conditions with 12-h light/12-h dark cycle at 22°C and 50% relative humidity. The female mice were randomly divided into two groups (n=15/group) according to their initial body weight. They received the modified gestating and growing formula (D10012G), containing 5000 (VD-C, Control group) or 25 (VD-D, VD deficient group) IU vitamin D3/kg diet for 6 week to obtain an optimal (>50nM) or deficient (<30nM) vitamin D status, respectively, in the serum; the VD levels were ascertained using five female mice before segregating them in the VD-C and VD-D groups. The other female mice (n=10/group) in each group were mated with 12-week-old C57BL/6J male mice by keeping two females per male in each cage. The mice were continuously fed throughout the gestation and lactation period.

On the postnatal day 21, serum was prepared from the blood collected from male pups born to mice in the VD-C and VD-D groups (n=5/group) and the contents of 25 hydroxyvitamin D3 (25(OH)D3) was determined. The other male pups were respectively fed high-fat (HFD, No. H10060, 34.9% fat by weight, 60% kcal) (n= 10/group, VD-C-HFD, VD-D-HFD) and normal fat (No. H10010, 4.3% fat by weight, 10% kcal) (n=10/group, VD-C, VD-D) diets with normal vitamin D content for 16 weeks, All the feeds were based on the formula of diet from Research Diets Inc. The schematic overview of the study design is shown in Supplementary Figure 1.

### Sample collection

After feeding for 16 weeks, blood samples were collected by retro-orbital bleeding from the male offspring fasted for 12h, and the animal were subsequently anesthetized by cervical dislocation to relieve them of their suffering. Serum was separated by centrifuging the blood samples at 3000 rpm for 15 min after allowing them to stand at room temperature (20-25°C) for 30 min. Immediately, after the sacrifice of the animals, the epididymal white adipose tissue (eWAT), inguinal white adipose tissue (iWAT), hepatic tissue, and pancreas were removed and weighted. Portions of these tissues were put in 10% buffered formalin for histological analysis, Some portions of eWAT and iWAT were stored in PBS for the analysis of immune cells. The remaining tissues were frozen in liquid N_2_ and transferred to a −80°C refrigerator until use. During the feeding process, the body weight was recorded weekly, and the food and energy intakes were measured monthly for the male offspring; the individual intake was calculated using the daily total intake for one cage divided by the number of mice in the cage.

To ensure the consistency of results, all the experiments were performed from 08:00 to 12:00 h in accordance with the recommendations in the Guide for the Care and Use of Laboratory Animals of National Administration Regulations on Laboratory Animals of China. The animal protocol was approved by the Committee on the Ethics of Animal Experiments of First Affiliated Hospital of PLA General Hospital in China.

### Biochemical analyses

The concentrations of calcium, triglycerides (TG) and total cholesterol (TC) in the serum were determined by colorimetric methods using the enzymatic assay kits (Maccura Biotechnology Co., Ltd, Sichuan, China). LC-MS/MS was used to measure the contents of 25(OH)D3 in the serum; this analysis was performed at the Analytical Facility for Bioactive Molecules of the China Food and Drug Administration (Beijing, China).

### Oral glucose and insulin tolerance tests

Oral glucose tolerance tests (OGTT) was performed in the 13th week of HFD feeding. After a 10-h overnight fasting and baseline sampling, mice were orally gavaged with 20% (weight/volume) glucose (2.0g /kg glucose solution) followed by collection of blood samples from the tail vein at 15, 30, 60, 90, and 120 min to determine the blood glucose levels using the Accu-Chek glucometer and glucose test paper (Johnson & Johnson, USA). Insulin tolerance tests (ITT) was performed 1 week after the OGTT. After 2-hour fasting, mice were weighted and blood samples were collected from the tail vein for serial blood glucose determinations. Biosynthetic human insulin (2IU/kg; Novo Nordisk A/S, Denmark) was injected through the intra-peritoneal route, and blood samples were collected from the tail vein at the baseline and after 15, 30, 60, 90, and 120 min of glucose challenge using Accu-Chek glucometer (Johnson & Johnson, USA).

### Histological analysis

Three samples from the four groups (VD-C, VD-D, VD-C-HFD, and VD-D-HFD) were randomly selected for histological analysis. Specifically, the hepatic tissues were stained with Oil red O for histopathological assessment and examined under a light microscope at 400× magnification. The eWAT and iWAT were embedded in paraffin and 6 μm sections were cut and stained with hematoxylin and eosin (HE). The stained sections were visualized under a light microscope at 200× magnification. The Image-pro Plus software was used to quantitatively analyze the size of fat cells in the eWAT and iWAT and the amount of lipid droplets in the hepatic tissues. All these analyses were performed by Servicebio, Beijing, China.

### Immunohistochemical determination of insulin in the pancreas and F4/80 in adipose tissues

The expression of insulin in the pancreas tissues obtained from VD-C, VD-D, VD-C-HFD and VD-D-HFD were assessed with rabbit anti-mouse insulin monoclonal antibody with slight modifications to account for segment-specific insulin distribution pattern. The F4/80 levels were determined immunohistochemically to estimate the quantity of the macrophages in the eWAT and iWAT. All these analyses were performed by Servicebio company in Beijing, China.

### Processing of fat pad for analysis of immune cells

The eWAT and iWAT stored in PBS were transferred to the RPMI medium. Collagenase II (Sigma-Aldrich Inc., St Louis, MO, USA) was added to the medium at a final concentration of 3mg/mL, and the suspension was incubated at 37°C for 45 min with constant shaking, and thereafter, filtered through a 200-mm membrane. The filtrate was centrifuged at 500×g/min for 8 min to pellet the the stromal vascular cells (SVCs); the floating adipocytes were removed by decanting the supernatant. The SVCs were incubated with 500μL red blood cell lysis buffer for 5 min at room temperature (20-25°C). The SVCs were resuspended in PBS containing 0.5% BSA prior to the incubation in flow cytometry block solution (Rat anti-mouse CD16/CD32, eBioscience, San Diego, CA, USA) for 15 min on the ice. Thereafter, the SVCs were stained with specific antibodies (CD45, CD8, CD3, CD4, CD11c, eBioscience; CD64, BD Pharmingen, Franklin Lakes, NJ, USA) for 30 min at 4 C in the dark. The cells were then washed twice with PBS and analyzed with the FACSCanto II Flow Cytometer (BD Biosciences, USA) using the FlowJo flow cytometry software (Treestar Inc., Ashland, OR, USA).

### RT-PCR assessment of the expressions of inflammatory cytokine genes

Total RNA in the eWAT and iWAT was extracted using the TRIzol Reagent (cat. no. 15596-206, Invitrogen, Carlsbad, CA, USA), and cDNA was reverse transcribed by SuperScriptTM III First-Strand Synthesis System for RT-PCR (cat. no. 18080-051, Invitrogen) following the manufacturer,s instructions. The expression levels of genes for pro-inflammatory cytokines, INOS, TNF-α, IFN-γ, IL-6 and IL-1β, in M1 macrophages and of those coding for anti-inflammatory cytokines, IL-4 and IL-10, in M2 macrophages were determined by RT-PCR (CFX-96, Bio-Rad, USA); the 36B4 genes was used as the invariant internal control. The oligonucleotide primers for the targeted genes were designed using Primer-BLAST (http://www.ncbi.nlm.nih.gov/tools/primerblast/) and are listed in Supplementary Table 1. The assays were performed in triplicates, and the results were normalized with respect the internal standard mRNA levels using the 2^−▲▲CT^ method.

### Determination of cytokine levels in the serum using Elisa kits

The levels of inflammatory cytokines, TNF-α, IFN-γ, IL-6, IL-4 and IL-10, in the serum were determined using Elisa kits, according to the manufacturer,s guidelines (Invitrogen, ThermoFisher Scientific, CN).

### Statistical analysis

All statistical analyses were conducted using SPSS 21.0. One-way analysis of variance (ANOVA) was performed to compare the means of indexes for different groups with normally distributed data, whereas the differences among data with non-normal distribution were assessed using Wilcoxon signed-rank test. The Student-Newman-Keuls (SNK) test was used to determine where the differences existed between two groups. A value of *P*<0.05 was considered to be statistically significant.

## Acknowledgements

This study was supported by National Natural Science Foundation of China (81602859). P.L., and P.L. participated the acquisition, analysis and interpretation of data. P.L., P.L., and L.L.Z. performed the mRNA extraction and gens expression. L.L.Z., and X.Y.C. carried out the animal feeding. Y.L.L., and W. J. L. assessed the plasma biochemical parameters. P.L., R.X.Z., K.M.Q., and Y.Z. drafted and revised the manuscript, which all authors have commented on. All authors read and approved the final manuscript before submitting.

